# Molecular mechanism of the dual regulation of bacterial iron sulfur cluster biogenesis by CyaY and IscX

**DOI:** 10.1101/212290

**Authors:** Salvatore Adinolfi, Rita Puglisi, Jason C. Crack, Clara Iannuzzi, Fabrizio Dal Piaz, Petr V. Konarev, Dmitri I. Svergun, Stephen Martin, Nick E. Le Brun, Annalisa Pastore

## Abstract

IscX (or YfhJ) is a protein of unknown function which takes part in the iron-sulfur cluster assembly machinery, a highly specialised and essential metabolic pathway. IscX binds to iron with low affinity and interacts with IscS, the desulfurase central to cluster assembly. Previous studies have suggested a competition between IscX and CyaY, the bacterial ortholog of frataxin, for the same binding surface of IscS. This competition could suggest a link between the two proteins with a functional significance. Using a hybrid approach, we show here that IscX is a modulator of the inhibitory properties of CyaY: by competing for the same site on IscS, the presence of IscX rescues the rates of enzymatic cluster formation which are inhibited by CyaY. The effect is stronger at low iron concentrations, whereas it becomes negligible at high iron concentrations. These results strongly suggest that iron-sulfur cluster assembly is an exquisite example of an enzymatic process which requires a double regulation under the control of iron as the effector.

## Introduction

Iron and sulfur are elements essential for life thanks to their unique redox properties. Yet, they are highly toxic. An efficient way to store them in cells in a nontoxic form is through formation of iron sulfur (Fe-S) clusters, labile prosthetic groups involved in several essential metabolic pathways (for a review (1, 2)). Assembly of Fe-S clusters is carried out by highly conserved machines which, in prokaryotes, are encoded by the suf, *nif* and *isc* operons. *Isc* is the most general machine with highly conserved orthologs in eukaryotes.

In *E. coli,* the *isc* operon contains eight genes (*i.e. iscR*, *iscS*, *IscU*, *iscA*, *fdx*, *hsca*, *hscb* and *iscX*) (3). Among the corresponding gene products, the most important players are the cysteine desulfurase IscS (<u>EC</u> <u>2.8.1.7</u>), which converts cysteine to alanine and IscS-bound persulfide (4), and IscU, a transient scaffold protein which forms a complex with IscS (5, 6). The last component of the machine according to the order of genes in the operon is IscX (also known as YfhJ), a small acidic protein about which very little is known (7, 8). IscX, the Cinderella of the *isc* operon, is not essential, in contrast to the other *isc* proteins (7). It is exclusively present in prokaryotes and in eukaryotes of the Apicomplexa, where it is highly conserved (9). Based on its phylogenetic occurrence, IscX seems to depend on the presence of IscS whereas the reverse is not the case (9).

The structure of *E. coli* IscX consists of a classical helix-turn-helix fold often found in transcription regulators (9, 10). *In vitro* studies have shown that IscX is able to bind IscS, thus suggesting a role for IscX as a molecular adaptor (9, 11, 12). IscX also binds to iron through a negatively charged surface, the same by which it recognises IscS (9). An exposed negatively charged iron binding surface which overlaps with the surface of interaction with IscS is also a feature present in another protein, CyaY (the ortholog in bacteria of the eukaryotic frataxin). This protein has attracted much attention because in humans it is associated with Friedreich’s ataxia (13). In contrast to IscX, frataxins are proteins highly conserved from bacteria to high eukaryotes (14) and are essential in eukaryotes (15). In prokaryotes, CyaY is external to the *isc* operon but has extensively been implicated in Fe-S cluster assembly (16). We have in the past shown that CyaY is an IscS regulator, which dictates the enzymatic assembly of Fe-S clusters (17, 18). Intriguingly, IscX and CyaY compete for the same site on IscS (11, 12, 18). A genetic interaction between CyaY and IscX was also demonstrated by a recent study which has validated a role of IscX as a new bona fide Fe-S cluster biogenesis factor(19). In some species, CyaY and IscX seem to replace each other.

These data raise the compelling question of whether there could be a functional link between CyaY and IscX, which could both elucidate the function of IscX and explain why these two proteins compete for the same binding site. Given the high conservation between prokaryotic and eukaryotic frataxins and desulfurases, answering this question could also inspire new studies on the regulation of eukaryotic frataxin. Using a complementary approach which makes use of enzymology, cross-linking and structural methods, we provide conclusive evidence indicating that IscX is a modulator of CyaY that switches off the inhibitory properties of CyaY as a function of the iron concentration. At low iron concentrations, IscS is under IscX control which has no inhibitory capacity. At high iron concentrations the system becomes controlled by CyaY. Based on our results, we suggest a general scheme that supports a role of CyaY as an iron sensor and provides the first testable indications concerning the function of IscX.

## Results

### IscX has different effects on the kinetics of cluster formation as a function of concentration

We started by exploring the effect of IscX on the rates of enzymatic assembly of the Fe-S cluster on IscU using an assay in which the cluster forms, under strict anaerobic conditions, through IscS-mediated conversion of cysteine to alanine and persulfide and is reconstructed on IscU (17). We performed the experiment at increasing concentrations of IscX in the range 1-50 μM using 1 μM IscS, 50 μM IscU, 250 μM Cys, 2 mM DTT and 25 μM Fe^2+^. We did not observe significant variations of the kinetics (Figure 1A,B) up to ca. 10 μM, a range in which the initial rates in the presence or absence of IscX were superposable, revealing that IscX does not have inhibitory effects under these conditions. Higher concentrations of IscX (from 20 to 50 μM, i.e. higher IscX/IscS ratios) led instead to inhibition of the reaction. Similar observations were made using circular dichroism (CD) (Figure 1C). The results at higher IscX/IscS ratios are in agreement with a previous study (12) in which the kinetics were performed at high concentrations of both Fe^2+^ (125 μM) and IscX (25 μM), but the results at lower IscX/IscS ratios are new and surprising. Why does IscS have a different behaviour at low and high ratios? There are different hypotheses which could explain these observations. First, the effect we observed at high IscX/IscS ratios could be explained by loss of available iron which could be sequestered by IscX, since we have reported elsewhere that IscX can undergo iron-promoted aggregation (9, 18). However, it was noted that aggregation occurs only when the assay is carried out at very low ionic strength. When salt is present, as was the case here, aggregation is not observed. Second, at low ratios the IscX occupancy on IscS could be too low to detect an effect because the complex has low affinity. Third, there could be a secondary binding site for IscX on IscS which is occupied only when the primary site is fully occupied. Only the 2:1 IscX-IscS complex would behave as an inhibitor.

**Figure 1.**
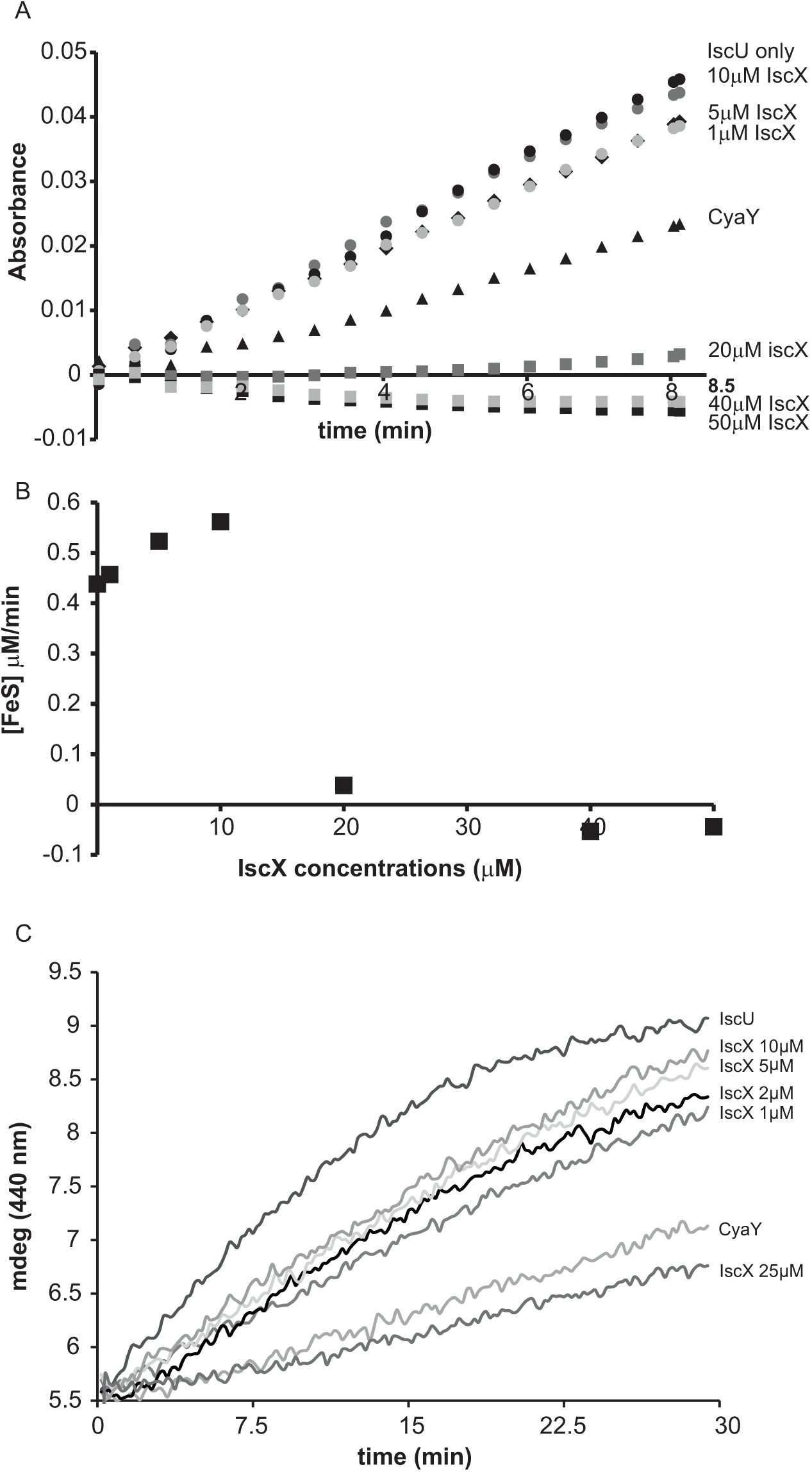
Effect of increasing concentrations of IscX on the enzymatic kinetics of Fe-S cluster formation on IscU. A) Enzymatic rates as measured by absorbance at room temperature. B) Plot of the initial rates at different concentrations of IscX. Increasing concentrations of IscX. Negative values are due to imperfect subtraction from the baseline and should be considered as zero (full inhibition). Only the initial part of the kinetics for each IscX concentration point was considered. The slope of these linear regions (for each IscX concentration) was used as the initial velocity. A molar extinction coefficient of 10.5 mM^-1^ cm^-1^ was assumed to convert the absorbance in concentration of Fe-S cluster produced. C) Kinetics followed by CD. The assays were carried out using 1 μM IscS, 50 μM IscU, 250 μM Cys, 2 mM DTT and 25 μM Fe^2+^ and increasing concentrations of IscX as indicated. For comparison, the experiment was also repeated in the presence of 5 μM CyaY. The error in repeated measurements was typically within 5% or less.

### The presence of IscU does not influence the affinity of IscX for IscS

To test the complex occupancy, we reconsidered the dissociation constants. We had previously estimated by fluorescence labelling and calorimetry dissociation constants (K_ds_) of 12 μM and 20 μM for the binary IscX-IscS and CyaY-IscS complexes respectively (9, 18). The IscX-IscS complex should thus be >50% populated under the conditions of our cluster formation assay. However, the presence of IscU could in principle modify these affinities as it is observed for binding to IscS when both IscU and CyaY are present. Also, although CyaY and IscX compete for the same site of IscS, the IscS dimer could have, at low IscX-IscS ratios, simultaneous occupancy of both CyaY and IscX. We used Biolayer Interferometry (BLI), a technique which can measure weak molecular interactions between several partners, to test this possibility. When we immobilized IscS on the surface and titrated with IscX only, we obtained a K_d_ of 8±3 μM for the IscX-IscS binary complex, in excellent agreement with the previous measurements (data not shown). When we immobilized CyaY on the surface, saturated with IscS and titrated with increasing quantities of IscX we obtained a K_d_ of 6±2 μM for the IscX-IscS complex (Figure 2A). Thus, the presence of CyaY on IscS does not modify the affinity of IscX for the desulfurase, and so major allosteric effects or binding cooperativity between the two proteins are unlikely. When we tested binding of IscX to the IscU-IscS complex (immobilizing IscS saturated with IscU), we obtained a K_d_ of 8±2 μM, a value comparable to the one obtained in the absence of IscU (Figure 2B). These data indicate that the presence of IscU bound to IscS does not affect the affinity for IscX and that the affinity of IscX for IscS is higher than that of CyaY. Thus, the effect observed at different IscX-IscS ratios cannot be ascribed to insufficient occupancy.

**Figure 2.**
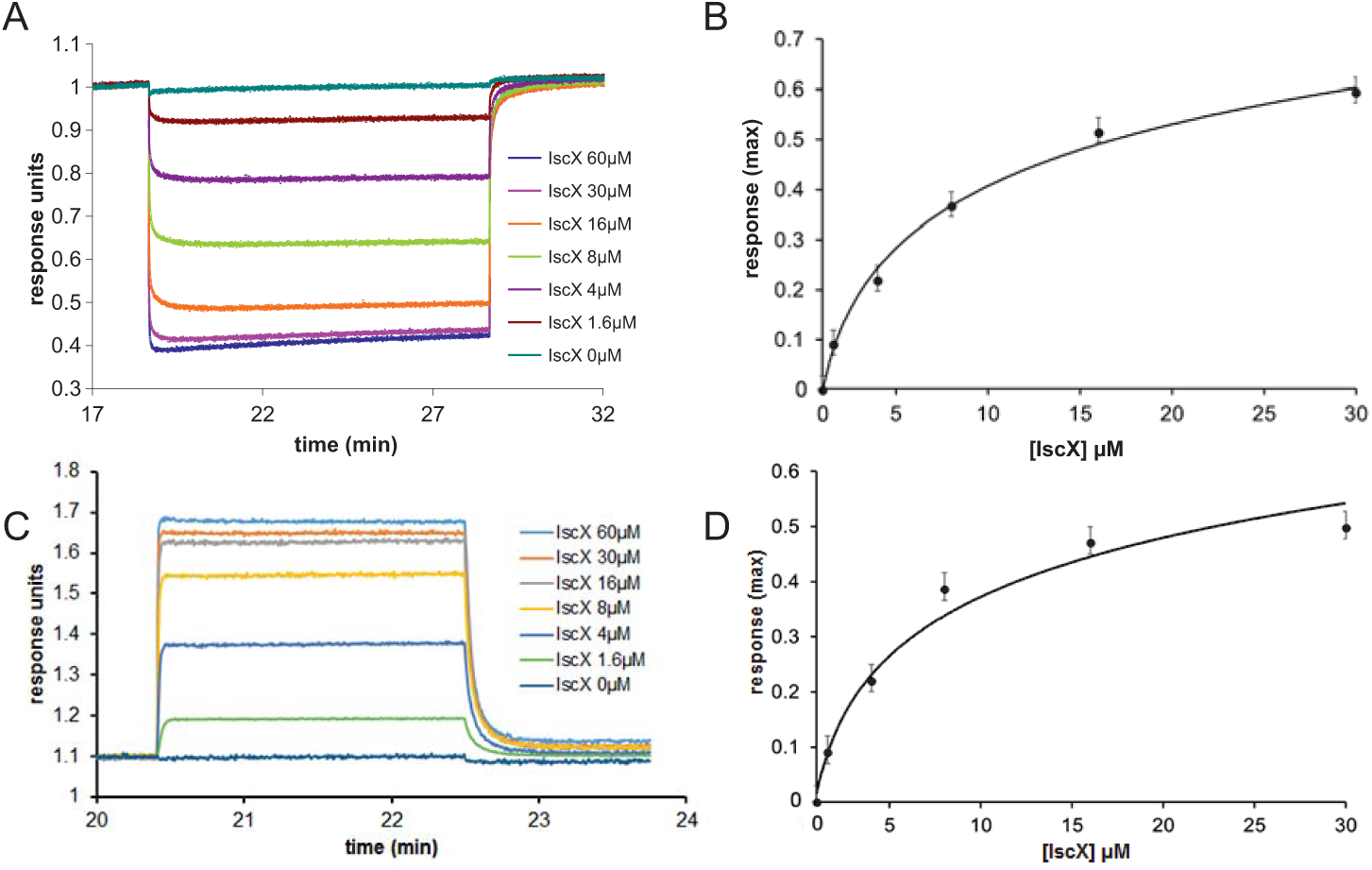
Quantification of the binding affinities of the IscX-IscS-IscU and IscX-IscS complexes to assess the influence of a third component. The experiment performed in tworked by completion. **A**) Plot of the interferometry response obtained by immobilizing CyaY saturated by IscS (10 µM) and titrating with increasing concentrations of IscX (0-60 µM). **B**) Plot of the response reduction as a function of IscX concentration. **C)** Plot of the interferometry response obtained by immobilizing IscS saturated by IscU (10 µM) and titrating with increasing concentrations of IscX (0-60 µM). **D**) Plot of the response as a function of IscX concentrations.

### IscX has two different binding sites on IscS

BLI proved inconclusive towards the presence of a secondary binding site. This could either be because we did not explore sufficiently high molar ratios, or because BLI depends on there being a significant change in the distance between the sensor's internal reference layer and the solvent interface. For formation of some complexes this can be rather small and thus undetectable. We thus used electrospray ionisation (ESI) mass spectrometry (MS) under non-denaturing conditions, a technique which, if optimised, provides direct information on all complexes present in a solution. The m/z spectrum (4100 – 6300 m/z) of IscS displayed well-resolved charge states (**Figure S1A**). The deconvoluted spectrum of IscS revealed a major peak at 91,035 Da consistent with the presence of dimeric IscS (predicted mass 91,037 Da) (**Figure S1B** and **Table 1**). Occasionally, shoulder peaks at +32 Da intervals on the high mass side of the peak were observed; these are likely due to the presence of one or more sulfane sulfur atoms, as previously observed for other proteins (20). An unknown adduct at +304 Da relative to the main IscS peak was also observed (**Figure S1B**). Addition of IscX to IscS to an 8:1 IscX/IscS molar ratio resulted in significant changes in the m/z spectrum. A complex pattern of new charge states, superimposed over those of IscS, were observed, consistent with the presence of IscX-IscS complexes (**Figure S2**). The deconvoluted spectrum (spanning 90 to 125 kDa) revealed the presence of four new species, corresponding to dimeric IscS in complex with up to 4 IscX molecules (at intervals of ~7.9 kDa) (Figure 3A). The observed mass of IscX was 7935 Da, +76 Da higher than the predicted mass of 7859 Da. LC-MS showed that a minor amount (<1%) of IscX was detected at the expected mass of 7859 Da, suggesting a covalent modification of the majority of the IscX protein. This is most likely due to adduct formation of IscX with *β*-mercaptoethanol during purification to give a mixed disulfide, which would have a predicted mass of 7935 Da (7859+78–2 = 7935 Da). Collision induced dissociation (CID) data supported this conclusion (**Figure S3**).

**Figure 3.**
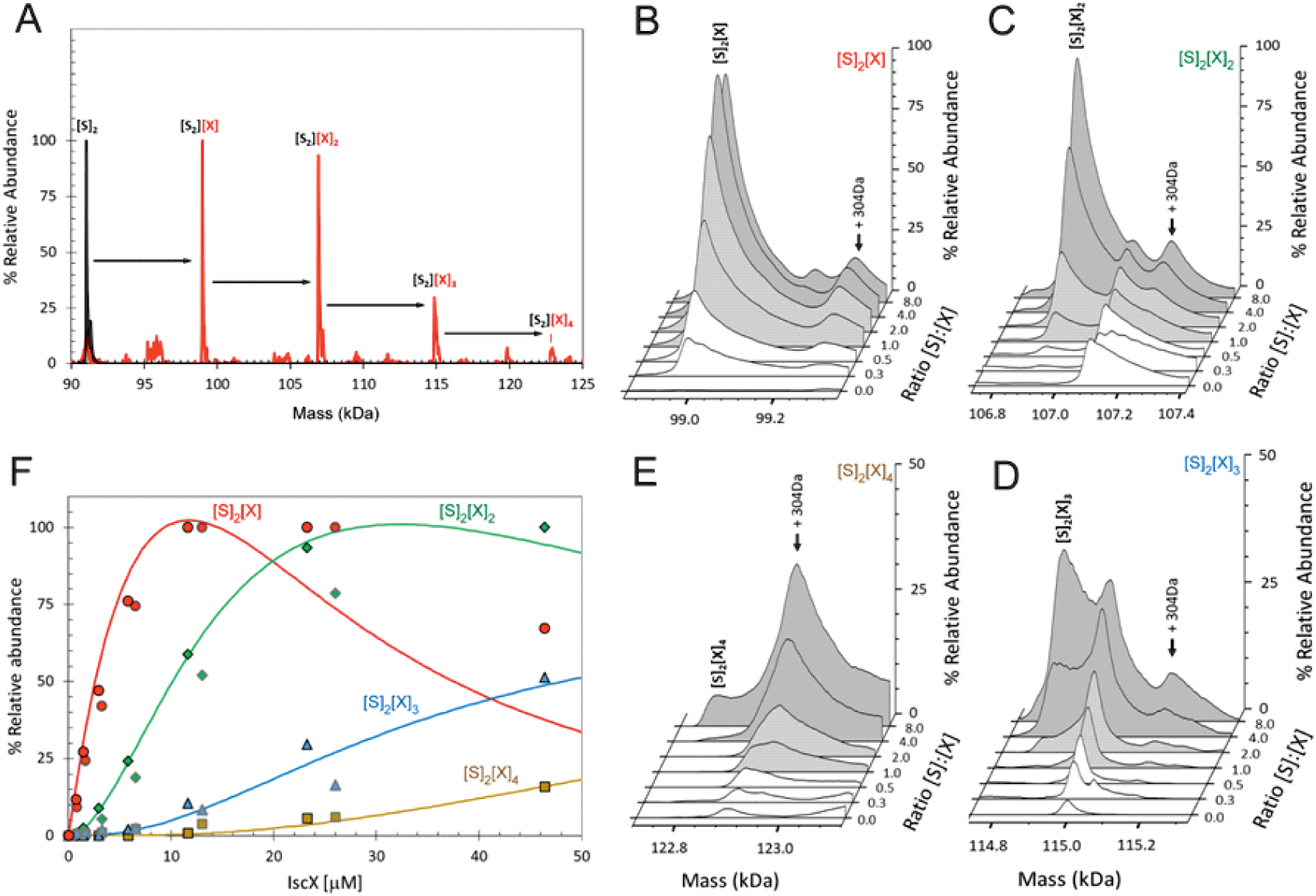
ESI-MS investigation of complex formation between IscS and IscX. (A) Deconvoluted mass spectrum of IscS over the mass range 90 – 125 kDa, showing the presence of the IscS dimer (black spectrum). Addition of IscX at an 8:1 excess gave rise to a series of IscX-IscS complexes in which the IscS dimer is bound by 1 – 4 IscX protein molecules (red spectrum). (B) – (E) Deconvoluted mass spectra at increasing ratios of IscX to IscS showing the formation/decay of the four IscX-IscS complexes, as indicated. A +340 Da adduct species is present in each of the spectra, including that of the IscS dimer, indicating that it originates from IscS. The precise nature of the adduct is unknown, but it is likely to arise from two β-mercaptoethanol heterodisulfides per IscS (4×76 = 304 Da), as the protein is in a solution containing 20 mM β-mercaptoethanol. (F) Plots of relative intensity of the four IscX-IscS complexes, as indicated, as a function of IscX concentration. Solid lines show fits of the data to a sequential binding model for 1 – 4 IscX per IscS dimer. IscS (3 µM) was in 250 mM ammonium acetate, pH 8. Note that abundances are reported relative to the most abundant species, which is arbitrarily set to 100%.

When increasing concentrations of IscX were added to IscS, we observed the gradual formation of dimeric IscS complexes containing 1 to 4 IscX monomers (Figure 3B-E). The complex of IscS with a single IscX protein, (IscX)(IscS)_2_, formed readily at low levels of IscX ([IscX]/[IscS]≈0.3) and maximized at [IscX]/[IscS]≈4 (Figure 3B). The (IscX)_2_(IscS)_2_ complex was detectable at a [IscX]/[IscS] ratio≈1 and reached maximum abundance by [IscS]/[IscX]≈8 (Figure 3C). (IscX)_3_(IscS)_2_ and (IscX)_4_(IscS)_2_ complexes were evident at [IscX]/[IscS] ratios ≈4 and ≈8, respectively (Figure 3D-F). The data were analysed according to a sequential binding model. The resulting fit of the data revealed that binding of the first two IscX molecules to form (IscX)(IscS)_2_ and (IscX)_2_(IscS)_2_ occurs with a similar affinity, *K*_d_=13.7±0.4 µM, consistent with the binding constant obtained by fluorescence and BLI. Binding of the third and fourth IscX molecules (to form (IscX)_3_(IscS)_2_ and (IscX)_4_(IscS)_2_) was also found to occur with comparable affinity, with a *K*_d_=170.0±10 µM, that significantly lower than the primary site. That the binding behaviour is well described by two pairs of dissociation constants indicates that the IscS dimer contains two equivalent pairs of IscX binding sites, with one primary (high affinity) and one secondary (low affinity) site per IscS protein.

This evidence conclusively supports the presence of two binding sites for IscX on IscS.

### Mapping the two binding sites by cross-linking

We next used cross-linking experiments to map the IscX binding sites on IscS, attempting to transform the non-covalent interaction in a stable covalent bond. In this assay, a cross-linking agent was added to link covalently proteins in close spatial proximity (21). We used bis[sulfosuccinimidyl]suberate (BS3), a cross-linking agent that reacts with primary amino groups up to ~11.5 Å apart, to mixtures of IscS and IscX at different molar ratios. The reaction produced two covalently attached protein complexes with apparent molecular weights of 52 kDa and 60 kDa, tentatively identified on PAGE as 1:1 and a 2:1 IscX-IscS complexes respectively (Figure 4). Mass spectrometry (MS) confirmed the presence of only these two proteins in both bands. Identification of the cross-link nature was achieved by subjecting to trypsin digestion the two species, as well as the isolated IscS. The obtained peptide mixtures were then analysed by MALDI/MS (**Table S2 of Suppl. Mat.**). Comparison between the spectra acquired for IscS and for the two complexes allowed us to detect a signal at *m/z* 3121.5±0.8 in the lower band which corresponds to the peptide 85-101 of IscS bound to the 1-9 region of IscX (theoretical mass 3120.6 Da). This value is uniquely compatible with a BS3-mediated cross-link involving Lys4 of IscX and Lys92, Lys93 or Lys95 of IscS. This identification was confirmed by HR LC-MS analyses and MS/MS data (**Figure S5 of Suppl. Mat.**). In the upper band, we could identify the same fragment together with another one (m/z 2424.2±0.5), which involves cross-linking between Lys101 of IscS and Lys4 of IscX (theoretical mass 2423.3 Da). As a control, we used an IscS mutant (IscS_R220/223/225E) which abolishes the interactions of IscS with IscX and CyaY (12, 18). We also used IscS_K101/105E which includes one of the lysines implicated in cross-linking with IscX. Formation of the IscX-IscS complex was strongly affected in both cases: binding was almost completely abolished with IscS_R220/223/225E, whereas the secondary binding site was barely detectable with the IscS_K101/105E mutant (Figure 4). As a control, when the same cross-linker was used with mixtures of CyaY and IscS only one band was observed (data not shown). These results map the positions of the two binding sites on IscS.

**Figure 4.**
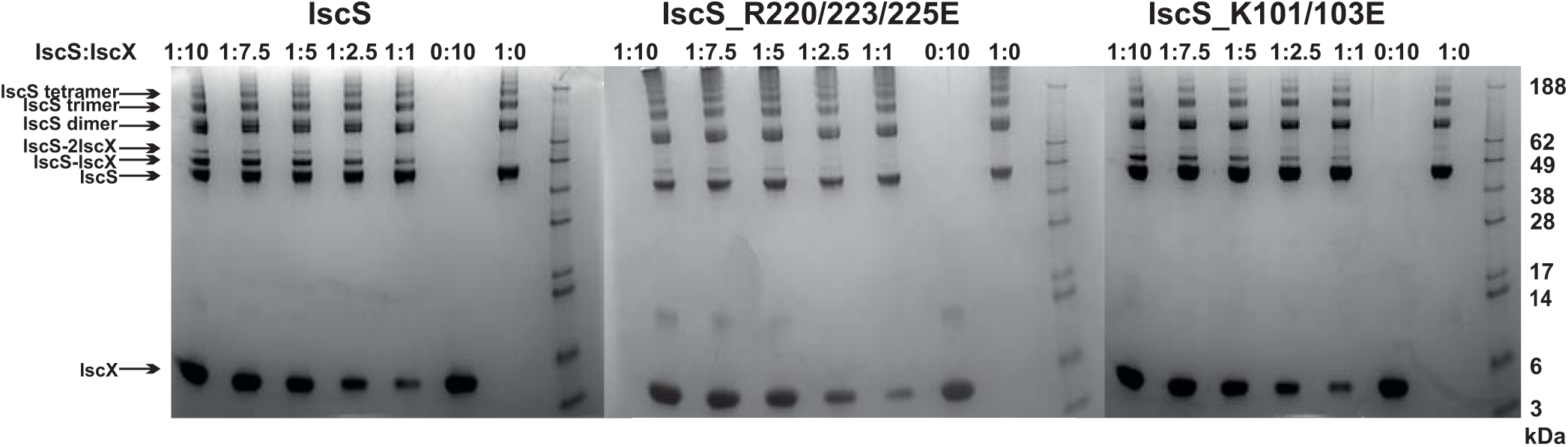
Cross-linking experiments to identify the interacting sites on IscX and IscS using different relative concentrations of IscS and IscX. A) Experiments using wild-type IscS (8 μM) and IscX at molar ratios 1:10, 1:7.5, 1:5, 1:2.5 and 1:1 μM from left to right. The last two lanes on the right are the controls carried out using isolated IscS and IscX. As expected the IscS dimer disassembles in SDS and runs as a monomer of ca. 45 kDa. B) The same as in A but with the IscS_K101/105E mutant. C) The same as in A but with the IscS_R220/223/225E mutant. The markers are indicated on the right.

### Structural characterization of the secondary binding site of IscX on IscS

Since it is difficult to assess the significance of these binding sites without a working structural model, we used small-angle X-ray scattering (SAXS) to grasp the shape of the 1:1 and 2:1 complexes. We recorded the scattering intensity patterns I(s) as a function of the momentum transfer *s* and the computed distance distribution function of IscX-IscS data (at 1:1 molar ratio, 5 mg/ml) (Figure 5A). The symmetric shape of the distance distribution function *p*(*r*) (Figure 5A, **inset**) is typical of a globular protein. The D_max_ value of the IscX-IscS complex (11.0 ± 0.5 nm) is comparable with that found for the CyaY-IscS complex and significantly lower than that observed for the IscU-IscS complex (12.1 nm). This is compatible with an arrangement in which IscX does not bind along the main axis of the IscS dimer but laterally increasing the IscS shape globularly [17]. The experimental Rg and excluded volume (V_Porod_) (3.09±0.04 nm and 135±10 nm^3^, respectively) are almost equal to the IscS homodimer (3.09±0.04 nm and 136±10 nm^3^, respectively) which suggests a partial dissociation of the complex (**Table S3 of Suppl. Mat.**). The molecular mass (92±5kDa) is also lower than that expected for a 1:2 IscX-IscS complex with full occupancy (117 kDa). This is in agreement with the presence of unbound species in solution due to the weak binding constants. The Rg and VPorod significantly drop when excesses of IscX were used (3.01± 0.03 nm/56±5nm^3^ for the 20:1 ratio and 2.75±0.03 nm/31±4nm^3^ for the 40:1 ratio). The higher the volume fraction of small species in the mixture (i.e. IscX molecules), the smaller is the average estimated excluded volume of the system. To further prove complex formation upon addition of IscX, all scattering curves recorded at different concentrations and different stoichiometries were analysed using a model-independent Singular Value Decomposition (SVD) approach (22). SVD clearly points to the presence of four components contributing to the scattering data in agreement with the presence of free IscS, free IscX, and their 1:1 and 2:1 complexes (**Figure S6 of Suppl. Mat.**).

**Figure 5.**
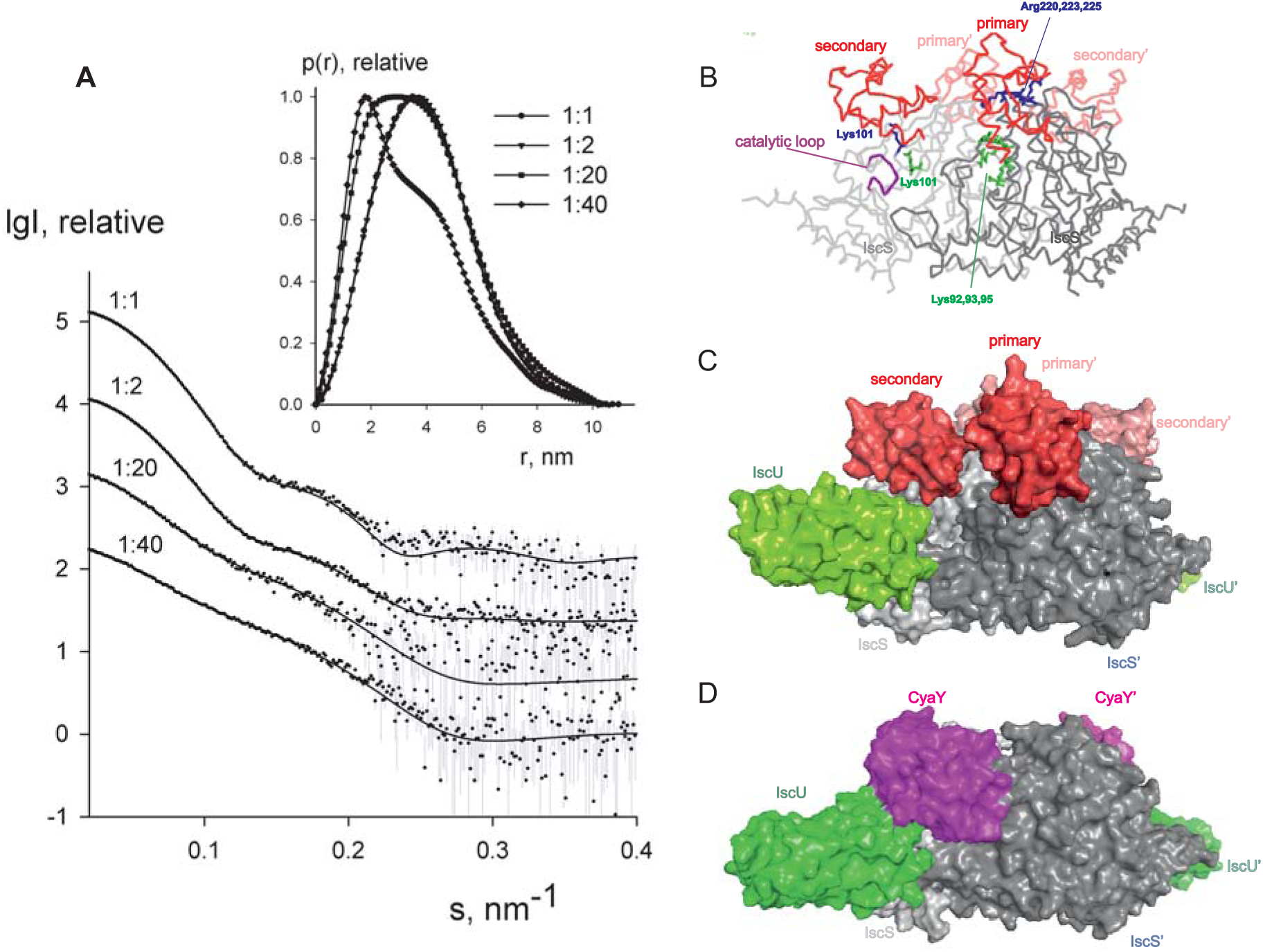
SAXS measurements. A) X-ray scattering patterns for IscX-IscS at molar ratios of 1:1 (at 5 mg/ml solute concentration), 2:1 (at 3 mg/ml), 20:1 and 40:1 (both at 0.5 mg/ml). The experimental data are displayed as dots with error bars. The scattering from a mixture of two rigid body models (1:1 and 2:1 IscX-IscS complexes) obtained by SASREFMX (for data with 1:1 molar ratio) and the fits by mixtures of unbound IscX, IscS dimers and 1:1/2:1 rigid body complexes (for data at 2:1, 20:1 and 40:1 molar ratios) as obtained by OLIGOMER are shown by solid lines. The plots display the logarithm of the scattering intensity as a function of the momentum transfer. The distance distribution functions are indicated in the insert. B) Backbone representation of the 2:1 IscX-IscS complex. The two IscS protomers are shown in different shades of grey. Residues which are either involved in cross-linking or have an effect on binding are indicated. The catalytic loop of IscS is indicated in purple. C) Same as in B) in a full atom representation. For reference, IscU (in green) is included (3LVL) to clarify the relative positions of the components. D) Comparison with the model previously obtained for the ternary complex CyaY-IscS-IscU [17]. Light and dark gray represent each of the protomers of the IscS dimer.

To construct a low resolution model of the IscX-IscS complex, rigid body modelling was used taking into account the possibility of 1:1 and 2:1 complexes. Multiple runs of SASREFMX starting from random initial configurations with applied P2 symmetry yielded models with qualitatively good fits to the data at a 1:1 molar ratio with a χ^2^ of 1.7. The interaction sites of IscS and IscX determined by NMR and cross-linking experiments were used as restraints (9, 12). The presence of unbound (free) components were taken into account by using free IscS dimers and IscX monomers as independent components in addition to the SASREFMX models. Their relative volume fractions were refined with the program OLIGOMER (**Table S4**). The volume fraction of free IscS decreases at a large excess of IscX consistent with an almost full occupancy of IscX. While it is difficult to distinguish between the fitting for 1:1 and 2:1 complexes, fits assuming both types of complexes are marginally better than those obtained assuming the presence of 2:1 complexes only. We estimated ca. 60-70% of 2:1 IscX-IscS complexes. The structure of the 1:1 complex is in excellent agreement with a previous model obtained by the same hybrid technique (12), IscX sits in the catalytic pocket of IscS contributed by both protomers of the IscS dimer (Figure 5B,C). This is the site where also CyaY binds (12, 18) (Figure 5D) thus explaining the direct competition between the two molecules. Experimental validation of this binding site has been supported by mutation studies of IscS(11). The second site is close but shifted towards the IscU-binding site.

The SAXS model suggests a logical explanation for the observed effects. We have previously demonstrated by molecular dynamics that binding of CyaY in the primary site restricts the motions of the catalytic loop which is thought to transfer persulfide, bound to Cys237, from the catalytic site to IscU (23). As compared to CyaY, IscX is smaller and would not be sufficient to block the movement of the catalytic loop as long as only the primary site of IscX is occupied. In this position, IscX simply prevents CyaY from binding, having higher affinity, but does not act as an inhibitor. However, binding of another IscX molecule with its N-terminus close to K101, as observed by cross-linking, can strongly increase the steric hindrance and result in inhibition of iron sulfur cluster formation resulting in an effect similar to that observed for CyaY.

### Understanding the competition between IscX and CyaY

We then explored more closely the conditions under which IscX competes with CyaY (11, 12). We first repeated the enzymatic assays keeping the conditions unchanged (1 μM IscS, 50 μM IscU, 250 μM Cys, 2 mM DTT and 25 μM Fe^2+^), adding CyaY (5 μM) and progressively increasing the concentration of IscX in the range 0-50 μM. Below 10 μM (i.e. below 2:1 IscX-CyaY and 10:1 IscX-IscS molar ratios), where IscX has no effect on IscS activity as demonstrated above, the experiment results in a progressive rescuing of CyaY inhibition (**Figure S7**), in agreement with a competition between the two proteins for the same binding site on IscS. The effect of IscX becomes noticeable at substoichiometric concentrations as compared to CyaY, as expected from the respective dissociation constants. However, even under the maximal condition of rescuing activity (10 μM), IscX is able to reach only ca. 70% of the rate compared to a control experiment performed in the absence of CyaY where there is no inhibition. This reflects the fact that, at relatively high IscX/IscS ratios, the secondary binding site, which also leads to inhibition, starts being populated probably favoured by a conformational change or an electrostatic rearrangement. Above a 10:1 IscX/IscS ratio, increase of the IscX concentration results in progressively marked inhibition (Figure 6A). When we titrated instead the system with CyaY (0-20 μM), keeping IscX fixed at 5 μM, we started observing a decrease of the intensities at 1:1 IscX/CyaY molar ratios and a marked inhibition for higher CyaY ratios (Figure 6B). We then put our results within the context of the protein concentrations observed in *E. coli*. Several independent studies have reported these under different growth conditions (24–27). Under non-stress conditions, both CyaY and IscX are usually, but not always, present at substoichiometric ratios as compared to IscS, with IscX in excess over CyaY (25). Different conditions can however remarkably change the molar ratios. We thus repeated the enzymatic assay using concentrations comparable to the non-stress conditions to verify that IscX controls the reaction under these conditions (4.1 μM IscS, 250 μM Cys, 2 mM DTT, 1.0 μM IscX, 0.7 μM CyaY and 25 μM Fe^2+^). We used IscU in a large excess (50 μM) because it is the reporter. We preferred to use this protein rather than another reporter because the IscU binding site on IscS does not interfere with CyaY or IscX (9, 18). At these ratios we do not observe inhibition indicating that the reaction is under the control of IscX (Figure 6C). Taken together, these results fully confirm the competing role of IscX and CyaY.

**Figure 6.**
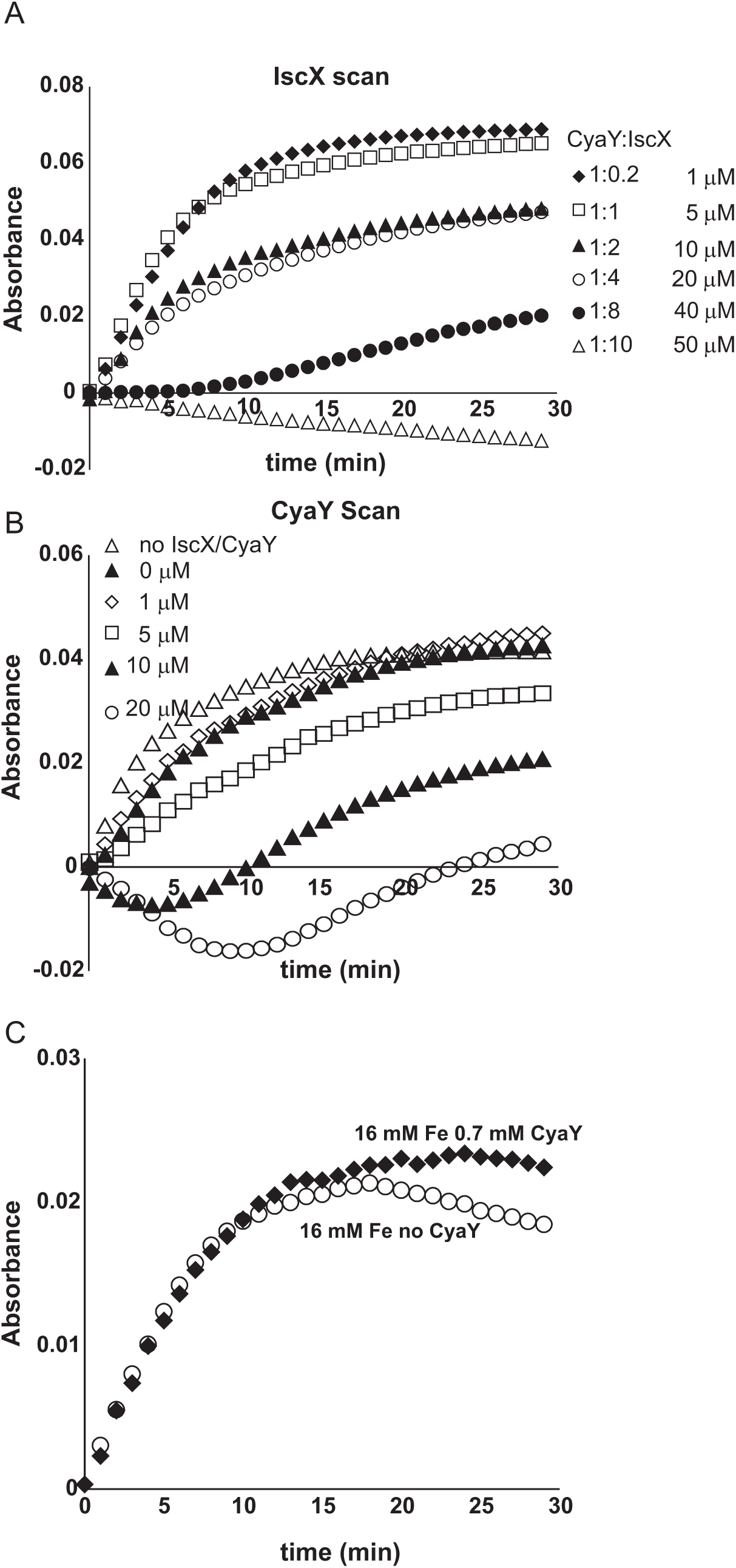
Competition between IscX and CyaY. A) Assay carried out using 25 μM Fe^2+^ in the co-presence of CyaY (5 μM) and increasing concentrations of IscX in the range 0-50 μM. When both CyaY and IscX are present, there is a competition which, under otherwise the same conditions, depends on the IscX:CyaY ratios. B) As in A) but with the concentration of IscX fixed to 5 μM and various concentrations of CyaY. C) Enzyme kinetics carried out with concentration ratios similar to those observed in the cell (4.1 μM IscS, 250 μM Cys, 2 mM DTT, 1.0 μM IscX, 0.7 μM CyaY and 25 μM Fe^2+^).

### The modulation effect of IscX and CyaY on cluster formation on IscU is iron dependent

Finally, we tested the effect of the competition between CyaY and IscX on Fe-S cluster formation on IscU as a function of increasing iron concentrations keeping the other concentrations unchanged (1 μM IscS, 50 μM IscU, 250 μM Cys, 2 mM DTT) because we had already observed that CyaY inhibition is enhanced by iron (17) (Figures 7A,B). As a control, we compared the variation of the initial rates as a function of iron concentration in the presence of IscU or CyaY individually. Cluster formation in the presence of the acceptor IscU only increased with the concentration of iron. The presence of CyaY (5 μM) drastically reduced the rate of cluster formation. The difference between the initial rates in the absence and in the presence of CyaY increased at increasing concentrations of iron. Addition of IscX to the reaction together with CyaY (1 and 5 μM respectively) generated a “rescuing” effect but only at low iron concentrations. Under these conditions, the effect reached a maximum around 15 μM iron. At higher iron concentrations, the curve obtained in the presence of both CyaY and IscX became similar to that obtained with CyaY alone. This implies that, at lower iron concentrations, IscX has higher affinity for IscS than does CyaY, impairing the ability of CyaY to inhibit cluster formation. The effect is reversed at higher iron concentrations where CyaY must acquire a higher affinity.

**Figure 7.**
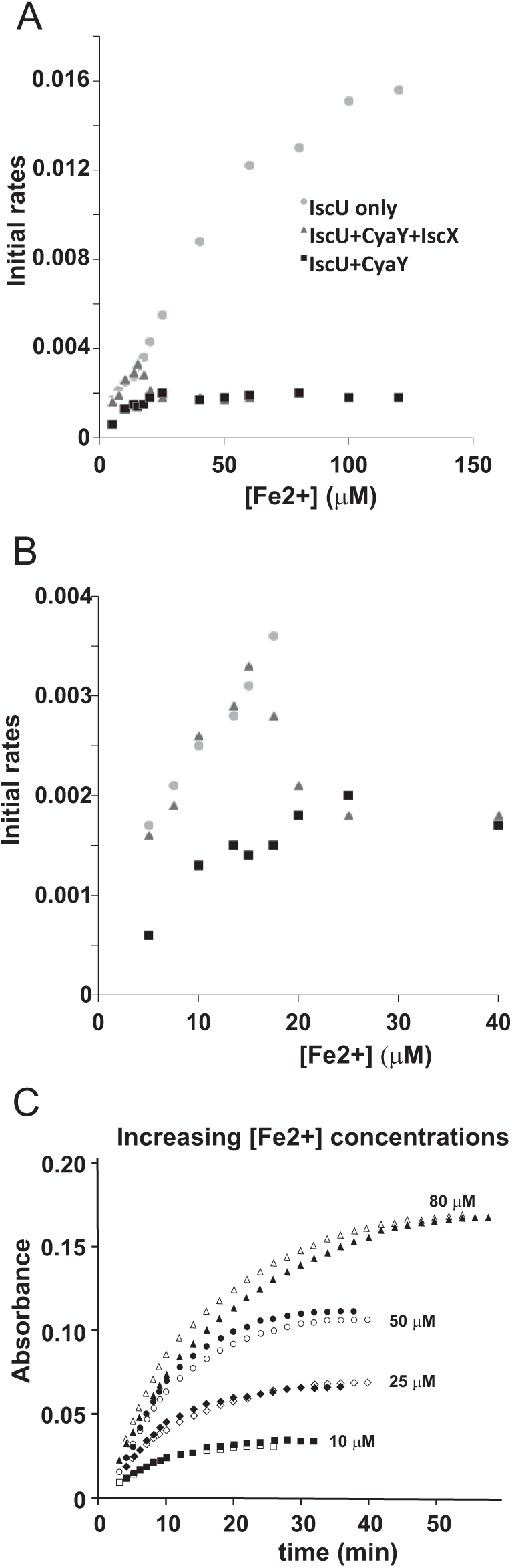
Iron dependence of the relative effect of CyaY and IscX on cluster assembly enzymatic kinetics on IscU. A) Initial rates of kinetics of cluster formation on IscU as a function of iron concentration in the presence of IscU only (grey rectangles), IscU and CyaY (black rectangles) and IscU, IscX and CyaY (grey triangles). B) The same as in A) but expanded for clarity. C) Assays carried out in the absence and in the presence of IscX (1 μM) at increasing concentrations of iron (10 μM, 25 μM, 50 μM, 80 μM) to assess whether IscX is affected by iron. No CyaY was added to the assay. All experiment contained 1 μM IscS, 50 μM IscU, 250 μM Cys, 2 mM DTT.

To dissect whether iron dependence is controlled by CyaY, IscX or both proteins together, we fixed the concentration of IscX to 1 μM and repeated the experiments varying the concentrations of Fe^2+^ from 10 to 100 μM in the presence and absence of IscX in the absence of CyaY. We observed that the initial rates increased at increasing Fe^2+^ concentrations but were independent of the presence of IscX (Figure 7C). Thus, the interaction of IscX to IscS is Fe^2+^ independent in this range of concentrations. This is at variance with CyaY whose interaction with IscS is clearly iron mediated, as previously demonstrated (17): the same experiment carried out with CyaY showed a marked difference in the absence and presence of CyaY which increases with iron.

We thus must conclude that IscX acts as a modulator of CyaY inhibition, resulting from the ability of CyaY to sense iron and bind to IscS more tightly at high iron concentrations.

## Discussion

Since its discovery (28), iron-sulfur cluster biogenesis has rapidly become an important topic of investigation both because it is a molecular machine at the very basis of life and because of its crucial role in an increasing number of human diseases (29). The study of this topic in the bacterial system is also specifically important given the important role that iron-sulfur cluster in bacterial infections (30). Much progress has been made in understanding this metabolic pathway but several questions have so far remained unanswered. Why does IscS bind different proteins using the same surface and how is recognition regulated? What is the role of IscX in the *isc* machine? What is the relationship between IscX and CyaY? We have now tentative answers to these questions.

Previous studies carried out at high IscX concentrations (>10 μM) led to the conclusion that IscX, like CyaY, is an inhibitor of cluster formation (12). We could reproduce these results but demonstrated that they strongly depend on the relative and absolute concentrations used *in vitro*. At high IscX-IscS ratios (i.e. >10:1), the inhibitory effect of IscX is the consequence of a secondary binding site on IscS which becomes populated only once the primary site is saturated. At low IscX-IscS ratios, IscX has no effect on cluster formation but efficiently competes with CyaY and modulates its strong inhibitory power also at substoichiometric ratios.

This is an important result which changes our perspective on IscX: this protein is not just “another frataxin-like protein” but the modulator of the inhibitory function of CyaY in bacteria; it is what silences CyaY. Our data are fully consistent with and explain in molecular terms a previous report that conclusively demonstrated a genetic interaction between CyaY and IscX and an additive positive effect on cluster maturation upon deletion of both genes (19).

We characterized the 2:1 IscX-IscS complex to obtain, albeit at low resolution, clues about the mechanism by which CyaY and IscX act as inhibitors. The two proteins compete for the same binding site on IscS but IscX is appreciably smaller than CyaY. In the complex with only the primary site occupied, IscX can thus allow free movement of the catalytic loop, which transports the persulfide from the active site to IscU (23). Occupancy of the secondary binding site would interfere instead with the loop movement by steric hindrance, thus producing an inhibitory effect. We also demonstrated that the effect of IscX on cluster formation depends on the iron concentration and that the iron sensor is CyaY and not IscX, whose effect on IscS is iron independent.

We can thus suggest a model based on these results which fully explains the role of both proteins (Figure 8). At low iron concentrations, cluster formation would be under the control of IscX and thus continue unperturbed. At higher concentrations, CyaY would have higher affinity for IscS and thus control the cluster formation rate. Additionally, it is likely that, under iron stress conditions, CyaY is up regulated (or IscS down regulated). Independent studies have indeed reported the concentrations of proteins in *E. coli* (24–27) under different growth conditions. Under normal conditions, IscX and CyaY are in substoichiometric ratios as compared to IscS with CyaY being also lower than, or equimolar, to IscX. The concentrations of the components can however vary considerably (up to 6-8 times) (24–27) according to the conditions, suggesting a tight regulation in which the ratios between the concentrations of IscS, IscX and CyaY are fine-tuned according to the cellular conditions. Such a system represents a sophisticated way to control the enzymatic properties of IscS: a tight regulation is required for maintaining the number of Fe-S clusters formed under tight control at all times, to match the number of apoproteins present. If this balance is lost, there will be too many “highly reactive” Fe-S clusters not being delivered which will be degraded.

**Figure 8.**
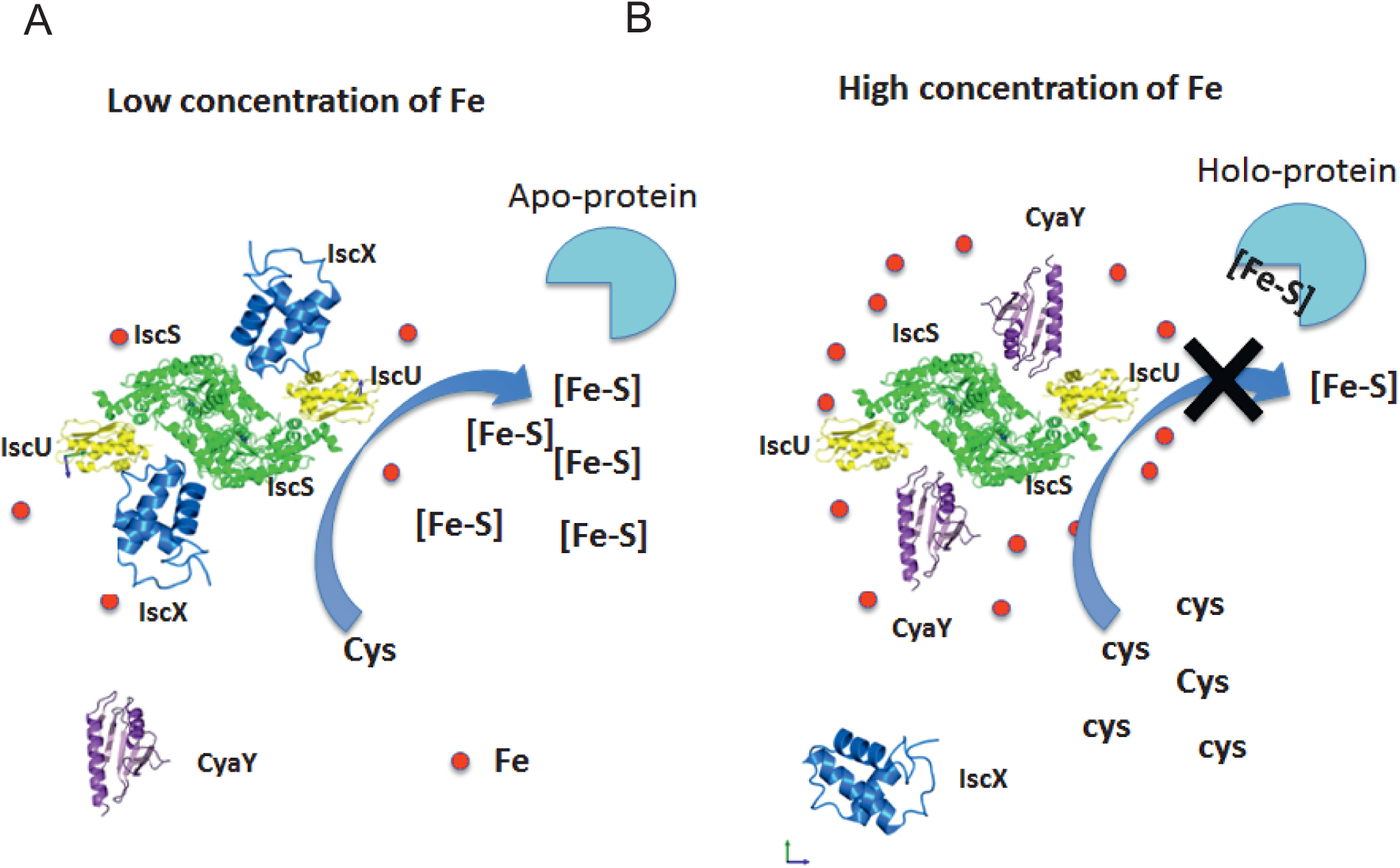
Model of the relative roles of CyaY and IscX. Left: at low iron concentrations, the system is under the control of IscX, which fills the binding site and impedes binding of CyaY. Right: at high iron concentrations, the system becomes under the control of CyaY.

From an evolutionary perspective, our results explain the presence of IscX in prokaryotes and not in most eukaryotes: bacterial IscS is a fully active enzyme, whereas the eukaryotic ortholog Nfs1 is inactive in the absence of its activators frataxin and the eukaryotic specific Isd11 and ACP (31, 32). As demonstrated in a recent paper, this behaviour can be explained by a different quaternary assembly of the desulfurase complex with Isd11 mediated by the acyl carrier protein in this structure (31, 32). These results indicate a different regulation of Fe-S cluster biogenesis in eukaryotes compared to prokaryotes. We propose the mechanistic basis of the bacterial regulation based on our results. We can also speculate that, as a regulator of the inhibitory function of CyaY in bacteria, IscX could have disappeared in the passage from prokaryotes to eukaryotes where the desulfurase was deactivated making the double repression unnecessary. This hypothesis would fully explain the difference in the function of frataxin between prokaryotes and eukaryotes, and suggests that IscX is the evolutionary “missing link”.

In conclusion, our results place the role of IscX into a new and completely different perspective and provide one of the most elegant examples of a double regulation of an enzyme in which inhibition and counter-inhibition are achieved by exploiting a triple-component system in which two proteins compete for the same binding site on an enzyme depending on the concentration levels of an effector, iron. Similar examples are not rare in nature and have been already described, for instance, in muscles where calcium is able to discriminate between different conformational states of tropomyosin and troponin I to determine different conformational states of myosin. Through our findings, we can account for the previous literature and suggest a new perspective for the quest to fully elucidate the roles of frataxin and IscX and their functions in the *isc* machine.

## Materials and Methods

### Protein production

All proteins used are from *E. coli*. Their sequences were subcloned in a pET 24d vector modified as fusion proteins with His-tagged glutathione-S-transferase (GST). The constructs were expressed in BL21(DE3). Bacteria expressing IscU were grown in Luria Broth enriched medium containing 8.3 MgZnSO_4_ (33) to stabilize its fold. The proteins were purified as previously described (9, 34) by affinity chromatography using Ni-NTA agarose gel and cleaved from the His-GST tag by Tobacco virus protease (TEV) protease. The mixture was reloaded on Ni-NTA gel to separate the His-tagged GST. This was further purified by gel-filtration chromatography on a Sephadex G-75 column. All the purification steps were performed in the presence of 20 mM β-mercaptoethanol. The protein purities were checked by SDS-PAGE and by mass-spectrometry of the final product. Protein concentrations were determined by absorbance at 280 nm using ε280 nm of 41370 M^-1^ cm^-1^ or 19480 M^-1^ cm^-1^ for IscS or IscX, respectively.

### Biolayer Interferometry

All experiments were performed in 20 mM HEPES (pH 7.5), 150 mM NaCl, 2 mM TCEP, 0.5 mg/mL BSA on an Octet Red instrument (ForteBio, Inc., Menlo Park, CA) operating at 25°C. Streptavidin coated biosensors with immobilised biotinylated IscS were exposed to different concentrations of IscX (0–50 µM). Alternatively, binding affinity between IscS and IscX was evaluated by competition with CyaY. For this experiment, streptavidin coated biosensors with immobilized biotinylated CyaY were exposed to 10 µM IscS at different concentrations of IscX (0-60 µM). Apparent K_d_ values were estimated by fitting the response intensity as a function of the concentration. Since BLI does not give information about reaction stoichiometry, the data were fitted to a simple 1:1 binding model using non-linear least squares methods (Levenberg Marquardt algorithm).

### Enzymatic assays

All enzymatic experiments to form Fe-S clusters on IscU were performed in an anaerobic chamber (Belle technology) under nitrogen atmosphere. The kinetics were followed at 456 nm as a function of time by absorbance spectroscopy using a Cary 50Bio Varian spectrophotometer. The initial rates were measured by incubating 1 μM IscS, 50 μM IscU, 250 μM Cys, 2 mM DTT and 25 μM Fe^2+^ in 50 mM Tris-HCl (pH 7.5) and 150 mM NaCl. When added CyaY was 5 μM. The reactions were initiated after half an hour incubation by addition of 1 μM IscS and 250 μM Cys. Each measurement was repeated at least five times on different batches of proteins. The data were always reconfirmed by CD performed under the same conditions.

### Cross linking assays

A mixture of 8 μM IscS was mixed with increasing molar ratios of IscX (0-80 μM) in PBS buffer. Bis[sulfosuccinimidyl]suberate was added to the protein to a final concentration of 2.5 mM. The reaction mixture was incubated at room temperature for 30 min and quenched by adding Tris-HCl (pH 8) to a final concentration of 20 mM. The quenching reaction was incubated at room temperature for 15 min. Identification of cross-linked peptides was performed using a classic mass-based peptide mapping approach (see Suppl. Mat). A further confirmation of the identity of the cross-linked peptides was achieved using the StravoX software, performing the “shuffle sequences but keep protease site” decoy analysis.

### Mass spectrometry under non-denaturing conditions

Solutions of IscS (3 µM) were mixed with increasing concentrations of IscX (0–16 IscX/IscS molar ratios). Samples were incubated at room temperature for 5 min before being loaded in a 500 µl gas-tight syringe (Hamilton) and infused directly in a Bruker micrOTOF-QIII mass spectrometer (Bruker Daltonics) operating in the positive ion mode. The ESI-TOF was calibrated online using ESI-L Low Concentration Tuning Mix (Agilent Technologies) and subsequently re-calibrated offline in the 4000 – 8000 m/z region. MS data were acquired over the m/z range 4,000–8,000 continuously for 10 min. For LC-MS, an aliquot of IscS or IscX was diluted with an aqueous mixture of 2% (v/v) acetonitrile, 0.1% (v/v) formic acid, and loaded onto a Proswift RP-1S column (4.6 x 50 mm, Thermo Scientific) attached to an Ultimate 3000 uHPLC system (Dionex, Leeds, UK). Processing and analysis of MS experimental data were carried out using Compass DataAnalysis v4.1 (Bruker Daltonik). Neutral mass spectra were generated using the ESI Compass v1.3 Maximum Entropy deconvolution algorithm. Titration data were fitted using the program DynaFit (BioKin, CA, USA). For further details see Suppl. Mat.

### SAXS measurements

SAXS data were collected on the EMBL P12 beamline on the storage ring PETRA III (DESY, Hamburg, Germany) (35). Solutions of IscX-IscS complexes were measured at 25°C in a concentration range 0.5-10.0 mg/mL at 1:1, 1:2, 1:20 and 1:40 molar ratios. The data were recorded using a 2M PILATUS detector (DECTRIS, Switzerland) at a sample-detector distance of 4.0 m and a wavelength of λ=0.124 nm, covering the range of momentum transfer 0.04<*s*<5.0 nm^-1^ (*s*=4*π* sin*θ*/*λ*, where 2*θ* is the scattering angle). No measurable radiation damage was detected by comparison of 20 successive time frames with 50 ms exposures. The data were averaged after normalization to the intensity of the transmitted beam and the scattering of the buffer was subtracted using PRIMUS (36). For further details on data treatment see Suppl. Mat.

## Acknowledgements

This work was supported by MRC (grant number U117584256) and BBSRC (grant BB/P006140/1).

